# Draft Genome of *Cochliopodium minus* (Amoebozoa): Insights into Its Complex Sexual Behavior, Across Domain Gene Acquisitions and Metazoan Type Signaling

**DOI:** 10.1101/2021.12.21.473654

**Authors:** Yonas I. Tekle, Fang Wang, Hanh Tran, T. Danielle Hayes, Joseph F. Ryan

## Abstract

To date, genomic analyses in amoebozoans have been mostly limited to model organisms or medically important lineages. Consequently, the vast diversity of Amoebozoa genomes remain unexplored. A draft genome of *Cochliopodium minus*, an amoeba characterized by extensive cellular and nuclear fusions, is presented. *C. minus* has been a subject of recent investigation for its unusual sexual behavior. *Cochliopodium’s* sexual activity occurs during vegetative stage making it an ideal model for studying sexual development, which is sorely lacking in the group. Here we generate a *C. minus* draft genome assembly. From this genome, we detect a substantial number of lateral gene transfer (LGT) instances from bacteria (15%), archaea (0.9%) and viruses (0.7%) the majority of which are detected in our transcriptome data. We identify the complete meiosis toolkit genes in the *C. minus* genome, as well as the absence of several key genes involved in plasmogamy and karyogamy. Comparative genomics of amoebozoans reveals variation in sexual mechanism exist in the group. Similar to complex eukaryotes, *C. minus* (some amoebae) possesses Tyrosine kinases and duplicate copies of *SPO11*. We report a first example of alternative splicing in a key meiosis gene and draw important insights on molecular mechanism of sex in *C. minus* using genomic and transcriptomic data.

## Introduction

Amoebozoa is a eukaryotic lineage that encompasses predominantly amoeboid organisms characterized by extreme diversity in morphology, ecology, behavior, and genomes^1^. Amoebozoa occurs globally in all major ecosystems, and is increasingly recognized as important in agriculture and ecology^2^. They are described from diverse natural habitats including marine, freshwater, and soil environments, and as symbionts or parasites affecting many livestock and humans. Amoeboid lineages are ecologically important as major bacteria grazers in fresh and marine environments. Some amoebas serve as a reservoir for life threatening human pathogens^3,4^. They also show complex life histories involving sexual and multicellular stages^5,6^.

Amoebozoa holds a key evolutionary position as one of the closest living relatives of Opisthokonta, a lineage that includes animals and fungi^7^. Despite their importance, the study of genomics in the group is at its infancy, limited to few lineages selected for their medical importance^8,9^ or lineages recognized as model organisms^10^. Despite insights gained from available amoebae genomes, the small number of sequenced amoebae represent only a small fraction of the phylogenetic breadth within the group. Comparative genome analysis of amoeboids covering diverse ecological and behavioral traits will enable investigation of many fundamental evolutionary questions including the evolution of life cycle histories, multicellularity, host-parasite co-evolution, amoeboid movement, and lateral gene transfer (LGT) across domains.

Recent studies in Amoebozoa using NGS techniques have generated large transcriptomic data that is contributing to our understanding of the group as a whole and making genome projects possible for less known amoebozoans such as *Cochliopodium*^11-14^. In this study, we sequence, assemble, and annotate a draft-level genome of an amoeba, *Cochliopodium minus*, a species characterized by extensive cellular and nuclear fusion^6,15^.

*Cochliopodium spp*. are lens-shaped, scale-bearing amoebae isolated from freshwater and marine habitats^15-17^. Some isolates, believed to be parasites, have been described from fish organs^18^. *Cochliopodium* is among the fast-evolving amoeboid lineages and its phylogenetic position within the supergroup has only been recently resolved using phylogenomic analysis^13,19^. *Cochliopodium* has been a subject of extensive study due to the unusual behavior of cell-to-cell interaction it exhibits in actively growing cultures during its life cycle^6,20,21^. *Cochliopodium* has long been considered asexual. However, our recent works demonstrate that this taxon engages both in cellular and nuclear fusion, followed by subsequent nuclear division and cell fission (plasmotomy), which are indicative of sexual activity^6^. The sexual nature of this behavior has been complemented using genetic and advanced cytological data^20,21^.

Sexual reproduction in most microbial eukaryotes is poorly documented including in the human pathogens (*Entamoeba, Acanthamoeba*) and in the amoeboid model organism (*Dictyostelium*). This is due in large part to challenges related to their complex and diverse quality of life cycles. For example, reported sex (meiosis) in most amoeboid microbes is assumed to occur during the dormant (cyst) stage^22,23^, which is a challenge for experimental study.

On the other hand, *Cochliopodium*, which has a well-documented life cycle^6^, displays sexual behavior during vegetative, active growth, stage, and therefore allows for overcoming century-old challenges that have stalled progress on elucidating sexuality in amoeboids. Sequencing the genome of *Cochliopodium minus* represents a key step in leveraging the advantages that this study system offers.

## Results

### Genome architecture and gene prediction of *C. minus*

We generated over 800 million sequencing reads (>100X coverage) using different sequencing technologies including 533.52 million Illumina short reads, 256,923 Oxford Nanopore MinION reads and 267 million 10X genomics reads. We obtained these genomic data from amplified DNA of single cells and nuclei pellets as well as from gDNA extracted from monoclonal cultures. We removed contamination from known food bacteria or associated entities, symbionts (e.g., viruses, archaea), and others from the environment following a series of bioinformatics and manual curation steps described in the methods. From these decontaminated data, we generated a draft genome of *C. minus* totaling 50.6 megabase pairs (Mbp) (Table 1). The assembly encompasses a total of 1,474 scaffolds, with average scaffold length of 34,311 base pairs (bps). The genome of *C. minus* is AT-rich with 26.33 % GC content (Table 1).

**Table 1.** Genomic composition and gene repertoire of *Cochliopodium minus* draft genome.

| Feature | *Cochliopodium minus* |
| --- | --- |
| Genome size (bp) | 50,573,891 |
| GC content (%) | 26.33 |
| DNA scaffolds | 1,474 |
| Longest scaffold length (bp) | 649,451 |
| Shortest scaffold length (bp) | 1,000 |
| Mean scaffold length (bp) | 34,311 |
| N50 (bp) | 126,531 |
| Total number of gene models | 19,925 |
| Proportion of genes with a size $\geq 300$ bp | 18,921 |
| Genes assigned to Cluster Orthologous Groups (COGs) | 12,256 |
| Non-ORFan genes | 14,202 |
| ORFan genes | 5,723 |
| Mean length of a coding gene (including introns) (bp) | 3,076 |
| Mean number of introns/genes | 4.7 |
| Mean number of exons/genes | 5.7 |
| Mean intron size (bp) | 682 |
| Mean exon size (bp) | 1,361 |

The overall genomic content including introns, exon and gene numbers are similar to most published amoebozoan genomes^8-10^. Our gene prediction was aided with transcriptome data of *C. minus*^11,20^ and a published genome of a closely related species, *Acanthamoeba castellanii*^9^. Using this approach, we generated a total of 19,925 gene models. Almost all transcripts obtained from the *C. minus* transcriptome were found in the draft genome with very high or full percentage matches, which is indicative of the quality and completeness of the assembled genome. In addition to this, 61.5% (12,256) of gene models were assigned to well-known biological processes in the Clusters of Orthologous Groups of proteins (COGs) database. The majority of gene models were classified under the Cellular Processes and Signaling (38.2%) category, followed by 19.8% and 15% under Metabolism, and Information, Storage and Processing categories, respectively (Table S1). The remaining 27.4% COG category included genes that are poorly characterized or of unknown function (Table S1).

Based on BLAST results, 28.7% of the 19,925 putative gene models constitute ORFans, genes with no BLAST hits to the NCBI GenBank database (Table 1). These gene models likely represent genes that have evolved extensively and are undetectable by BLAST as well as a subset of genes unique to *C. minus*. Among the ORFans, 98 and 41 were upregulated in fused and unfused cells of *C. minus*, respectively (Table S2). The fused amoeba cells have been shown to be associated with the sexual stage in the life cycle of *C. minus*^20^. While the exact functions of ORFans are unknown, a preliminary functional exploration of these upregulated genes using InterPro has revealed some common domains of proteins involved in different biological processes. Among the common domains found in the up-regulated ORFans in fused cells include those that are involved in protein–protein interactions (e.g. Fox-box, Ankyrin repeats, WWE), catalytic sites (AAA, Protein kinase), cytoskeletal proteins and signal transduction (e.g. RhoGAP, Calponin homology, CUE), DNA damage sensing (BRCT, PH), membrane-associated proteins (GRAM domain), transcription regulation (IBR domain, a half RING-finger domain), nuclear and replication associated (e.g. ORC, YabA) and Zinc finger motifs (Table S2).

### Taxonomic distribution of *C. minus* gene models

Similar to other amoebae genomes, the taxonomic distribution of gene models in the genome of *C. minus* show mosaicism of various taxonomic groups. The majority (54%) of the gene models matched eukaryotic genes. A substantial number of the gene models, ∼29%, are ORFans (Fig. 1). Among other living domains, the largest proportion (15% - 2995 genes) show highest similarities to bacteria, while only a small fraction shows highest similarities to archaeal genes (0.9%, 170 genes) (Fig. 1). An even smaller percentage of gene models (0.7%, 140 genes) show highest similarity to viral genes (Fig. 1). This latter set, non-eukaryote matching genes, makes up core components of cellular (signaling and metabolism) and information storage and processing (Fig. S1). Expression of some of these gene models with high similarity to bacterial, archaeal and viruses have been detected in transcriptome data sampled from various stages of the *C. minus* life cycle (Table S3).

**Figure 1.**
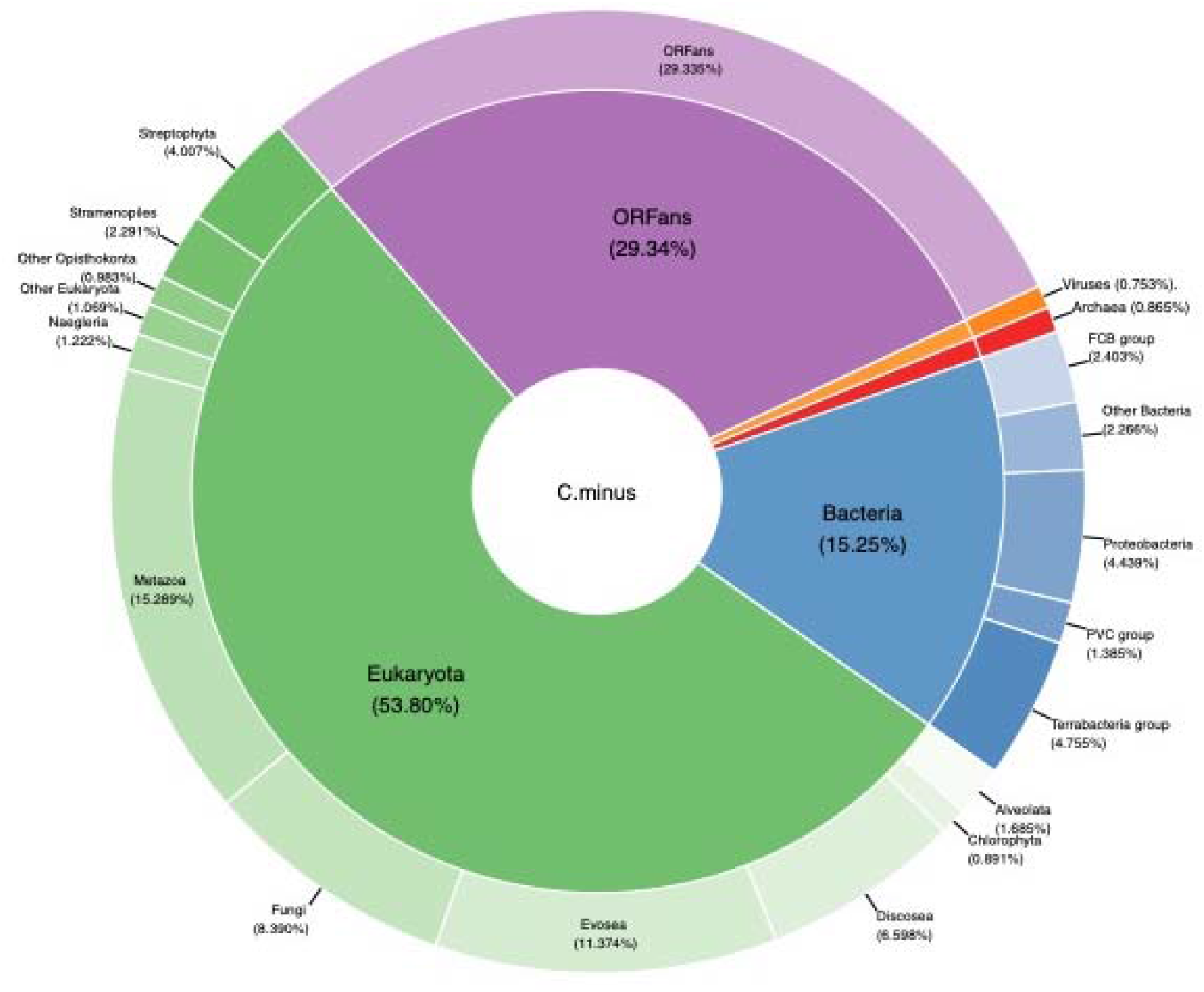
Taxonomic classification of predicted proteins deduced from *Cochliopodium minus* draft genome.

### Evidence of interdomain LGT in the *C. minus* draft genome

We used a combination of approaches to identify genes that are potentially acquired into the genome of *C. minus* through lateral gene transfer (LGT). Based on BLAST similarities and alien index analyses, we identified a large number of LGT candidates from bacteria and archaea. Out of the 2,995 gene models with bacterial origin (best homology matches with bacterial genes), 303 genes were shown to have alien indices above the default threshold (>45) indicating that they are putative LGT candidates (Table S3). Phylogenetic analysis of selected putative LGTs showed that these genes most closely resemble those from a range of bacterial taxa including novel bacterial (Candidatus) phyla (Fig. 2A, B, Table. S3). Included among the bacterial phyla that putatively represent origins of transferred genes are Proteobacteria (106), Terrabacteria group (65) and FCB group (56; Table. S3). Some of the putative LGTs found in the *C. minus* genome are also shared with other amoebae and eukaryotes (see Fig. 2B). While all of our selected bacterial-like genes and putative LGTs are found in scaffolds containing most amoebozoan and eukaryotic genes, in some rare instances, we observed an amoeba-like gene within contaminant scaffolds. It is likely that these genes might represent LGTs from amoebae to bacteria, however, we have not found strong evidence to suggest this is the case. It is also likely that this can be an assembly problem. In this study, only bacterial genes that straddle scaffolds dominated by amoeba and eukaryotic genes are considered in the final assembly of our draft genome. The observation of amoeba-like genes in bacterial scaffolds requires further investigation.

**Figure 2.**
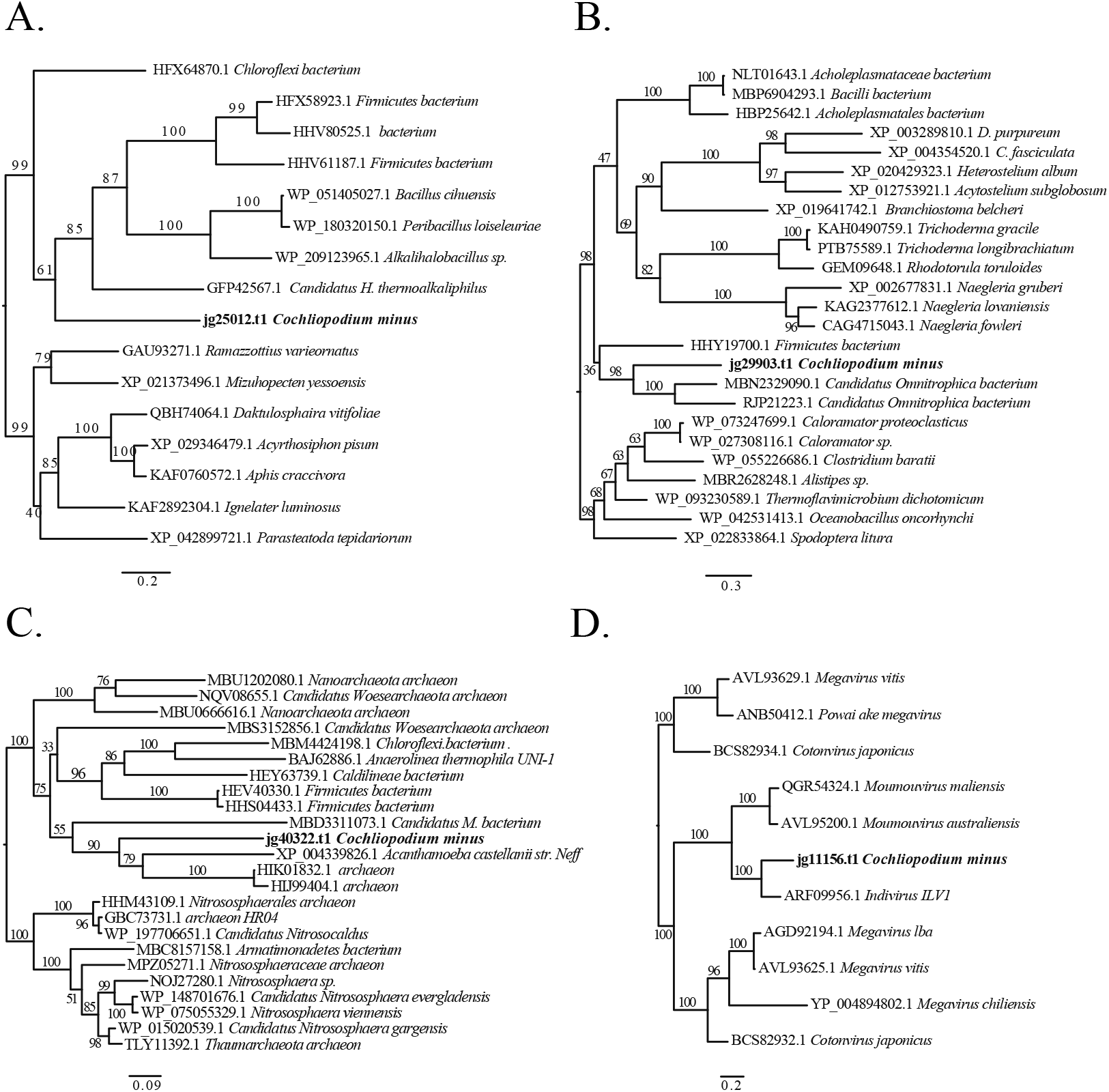
Phylogenetic reconstructions demonstrating putative lateral gene transfers (LGTs) in *C. minus* genome among bacteria (a, b), archaea (c), and giant viruses (d). Clade supports at nodes are ML IQ-TREE 1000 ultrafast bootstrap values. All branches are drawn to scale.

Among the 170-archaeal origin genes found in the *C. minus* draft genome, 10% (17 genes) of them had an alien index above the threshold (Table S3). Amoebae genomes contain much fewer genes of archaeal origin compared to bacterial genes and only few genes are reported to have been acquired laterally^9,24^. A phylogenetic analysis of the putative archaeal LGT showed that both *C. minus* and *A. castellanii* acquired a similar gene that encodes for Inosine-5’-monophosphate dehydrogenase (IMPDH) (Fig. 2D). The likely donors of the putative LGTs in archaea come from diverse taxonomic groups including Thaumarchaeota, Euryarchaeota, Thermococci and some Candidatus archaeon phyla (Table S3).

### Giant- and other-viruses origin genes in the genome of *C. minus*

The association of viruses with free-living amoebae and the incorporation of viral genes into the genomes of amoebae are well established^9,25,26^. Similar to other amoebae, the genome of *C. minus* carries genes of viral origin. A total of 140 gene models showed best homology matches with viral genes (Fig. 1, Table S4). As expected, the majority (90%) these genes have giant virus origin (Table S4), while the remaining includes double-stranded DNA (dsDNA) bacteriophages and unclassified viruses (Fig. 3). Among the giant virus, Pithovirus (33.3%) appears to be the dominant contributor of viral genes in the *C. minus* genome (Fig. 3). Several other genes also seem to originate from common giant viruses associated with free-living amoebae, which include Mimivirus, Fadolivirus, Indivirus, Moumouvirus, Pandoravirus and Marseillevirus (Fig. 3). A total of 49 genes (35%) have alien indexes above the threshold suggesting that they are likely candidates for LGT between these diverse viruses and *C. minus* (Table S3). Members of Mimivirus and Pithovirus are among the largest putative LGT donors in the *C. minus* genome (Table S4). A phylogenetic reconstruction of one of these putative viral LGT, collagen triple helix repeat containing protein, showing a close affinity to Indivirus is shown in Figure 2D.

**Figure 3.**
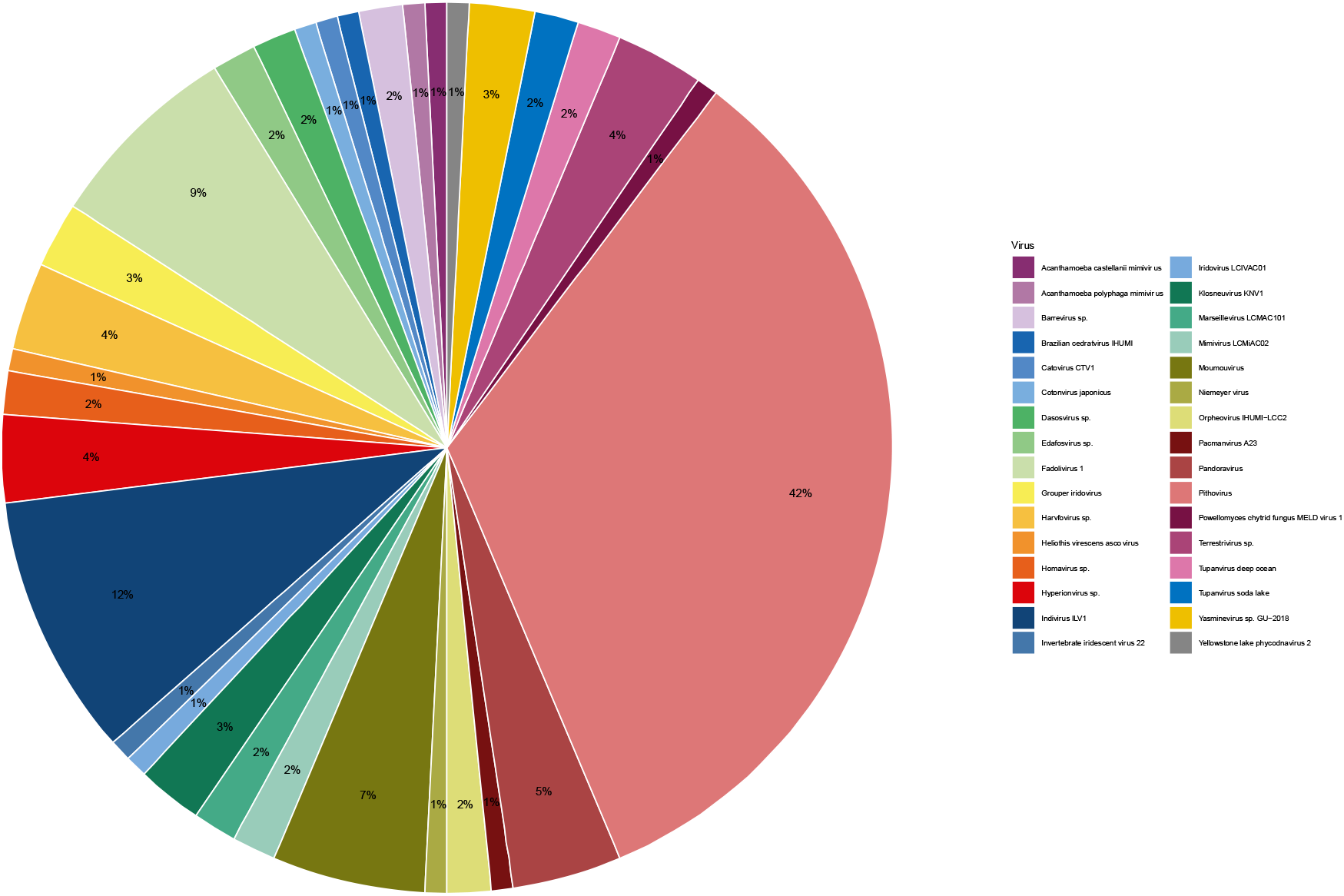
Taxonomic distribution of sequences matching to giant viruses in the *Cochliopodium minus* draft genome.

### Genes involved in sexual life cycle of *C. minus*

#### Meiosis genes

Most of the evidence for sexual reproduction in microbial eukaryotes comes from detection of genes involved in sex particularly those that are meiosis specific^27^. This is primarily because observation for direct (physical) evidence of sex is lacking for most microbial eukaryotes due to their complex and diverse life cycles^28^. Due to limited genome data, most of the genetic evidence for sex in Amoebozoa was deduced from transcriptome data^29^. While analyses of transcriptome data demonstrated the sexual nature of amoebozoans, the detection of the full complement of meiosis genes in such studies are sporadic due to the incomplete nature of the transcriptome data. Our previous study using this approach conclusively detected only four (*DMC1, HOP*2, *MND1, MSH4*) of the common meiosis specific genes inventoried in *C. minus* transcriptome^21^. Some genes that were detected such as *SPO11* were partial sequence, and their homology was questionable. Our thorough analysis of the draft genome of *C. minus* recovered almost the full complement of meiosis genes and revealed interesting information about the nature of the gene products. In addition to previously detected meiosis specific genes, we found *MSH5, MER3, REC8, SPO11, PCH2* and *ZIP4* in the draft genome of *C. minus*. Some of these genes (e.g., *SPO11*) appear to have at least one paralog. We find two copies (paralogs) of *SPO11* in draft genome of *C. minus* similar to *Acanthamoeba castellanii*^30^. These paralogs are identified as *SPO11-1* and *SPO11-2* based on phylogenetic analysis that included diverse eukaryotic groups (Fig. 4). Our comparisons of these *SPO11* paralogs to other amoebae revealed that some members of Amoebozoa likely possess (express) these two copies (Fig. S2). Similarly, we identified two copies of *MER-*like genes that are shown to group separately in a phylogenetic analysis in *C. minus* genome (Fig. S3). These paralogs possess duplicate copies of three domains (DEAD, Helicase and SEC63) significantly (>48%) differing in their primary sequence homology at nucleotide level (Fig. 5A). A closer examination of these three domains that make up *MER3*, showed that one of the genomic copies (jg36575.t1) shows a higher homology similarity to genes identified as *MER3* in human and other protists, while the other copy is likely a closely related gene (e.g., U5 snRNP) with similar structural domains (Fig. S3). The three domains in these paralogs are arranged in consecutive triplets (Fig. 5B). Comparison of the expressed transcript and the genome copy (jg36575.t1) of *MER3* showed that this gene undergoes post transcription processing. The composition of the processed transcript (jg36575.t1) includes the first two copies of domains (DEAD and Helicase) in conjunction with a second copy of the SEC63 domain (Fig. 5B). The first copy of SEC63 domain and the second copies of DEAD and Helicase domains appear to have been spliced out during pre-mRNA processing.

**Figure 4.**
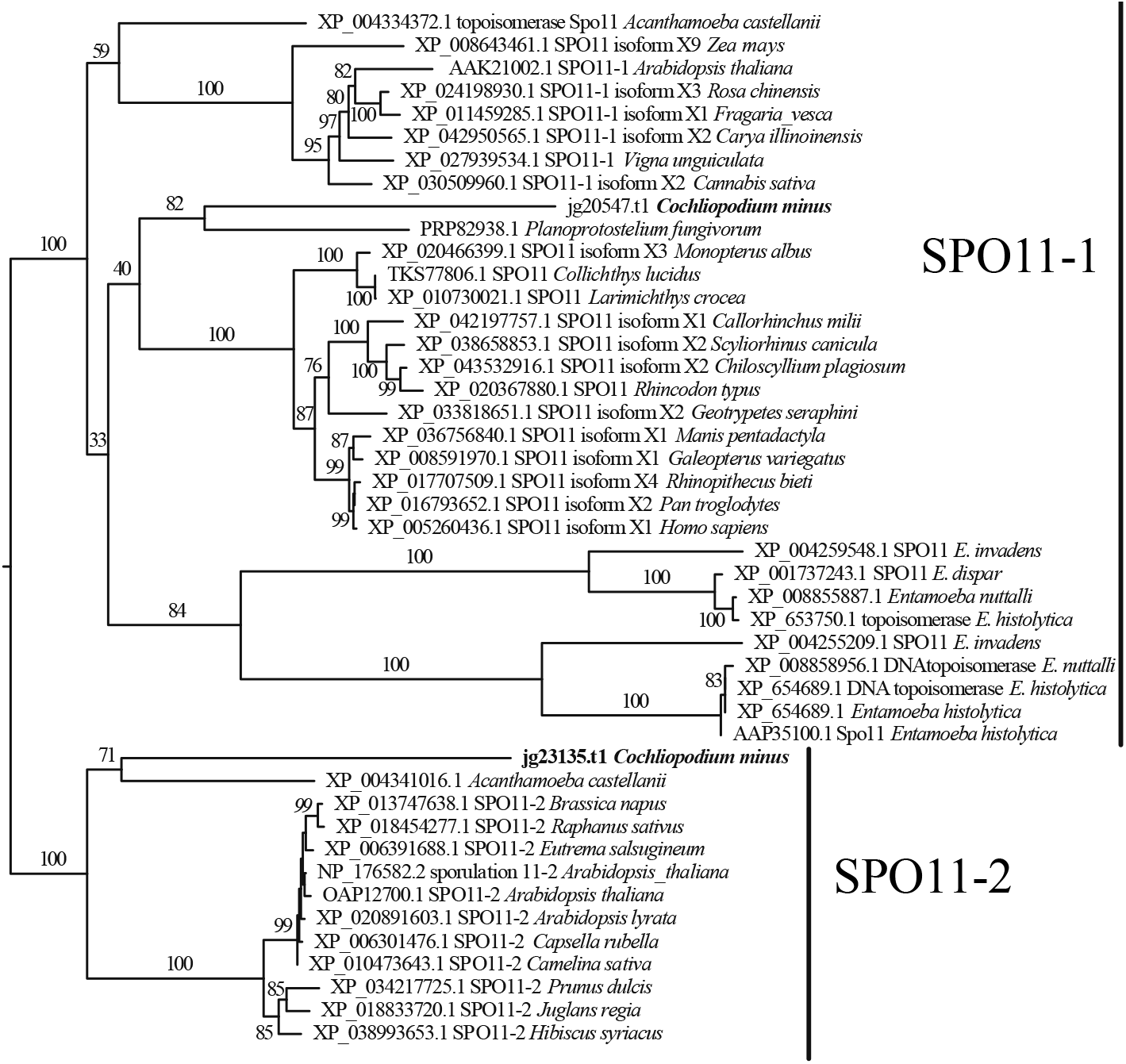
Phylogenetic reconstructions of *SPO11* paralogs. Two copies of *SPO11* genes from the *Cochliopodium minus* draft genome are placed into two well-supported clades. Clade supports at nodes are ML IQ-TREE 1000 ultrafast bootstrap values. All branches are drawn to scale.

**Figure 5.**
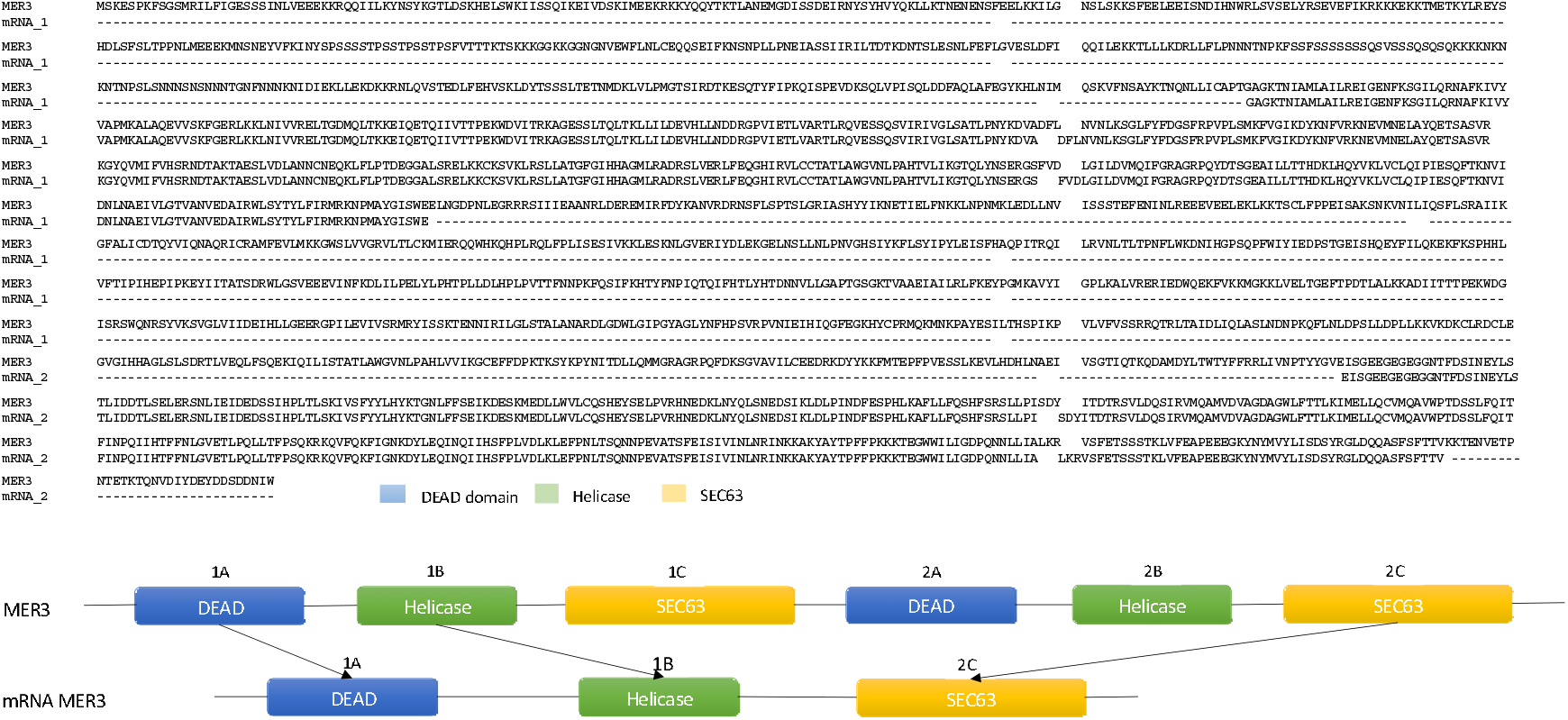
Alignment of *MER3* genomic and transcript (mRNA) copies and an illustration of alternative splicing resulting in mRNA *MER3* with three domains combination. Corresponding domain colors are used in alignment and illustration.

#### Cellular (plasmogamy) and nuclear (karyogamy) fusion genes

In a previous study, we searched *C. minus* transcriptome data for more than 30 genes known to be involved in cellular and nuclear fusion found in diverse eukaryotes^21^. We found only a few of these genes in these data including the plasmogamy genes: *BNI1, KEX2* and *MYO2* and the karyogamy genes: *CDC4, CDC28, CIN4, KAR3, KEM1* and *KAR2*. In our current analyses, we have found two additional plasmogamy genes: *CD9* and *RVS161* and 8 additional karyogamy genes: *RVS161, BIK1a, CDC34, CIN2a, KAR4, JEM1a, SEC63* and *SEC72a*. The identification of these genes was based on sequence homology, phylogenetic analyses, and domain analysis. We were unable to identify the fusogene, *HAP2* nor the karyogamy gene *GEX1/KAR5*. This was unexpected as these genes are commonly found in diverse eukaryotes including some amoebozoans^31^.

## Discussion

### Genome diversity of Amoebozoa and its significance

Genomic studies in Amoebozoa are contributing to our understanding of the evolution and origin of the supergroup as well as help answering fundamental questions pertaining to innovations and shared eukaryote cell features^8-10^. Previous genome studies have revealed that members of Amoebozoa possess cellular processes only thought to exist in complex eukaryotes. For example, Tyrosine kinases were thought to be the hallmark of metazoan and choanoflagellate evolution^32^. Genomic studies of amoebozoans revealed that some members (e.g., *Physarum polycephalum, Acanthamoeba castellanii, Entamoeba histolytica*) possess complete tyrosine kinase signaling toolkit similar to that of metazoans^9,33^. *C. minus* also possesses this complex signaling system (Table S5). This finding and the discovery of Tyrosine kinases outside of Metazoa reinforces the ancestral nature of Tyrosine kinases in eukaryotes albeit with multiple loses in some major lineages^33,34^. Genomic studies are also revealing variations, loses and innovation that reflect the great diversity of Amoebozoa. For example, *Dictyostelium discoideum* lacks a key gene that is used in the initiation of meiosis (see below). Similarly, *D. discoideum* lacks tyrosine kinase gene toolkit found in other amoebae but possesses alternative genes that play a similar role^35^. More genomic studies will unravel the diversity and evolution of signaling in amoebozoans.

Among sequenced genomes of amoebozoans with annotation (NCBI accessed December 7, 2021), the genome size of *C. minus* falls among the largest (Table S6). Most of the amoebae genomes sequenced to date are very small sized (average size 33 MB) relative to what is reported of the supergroup that includes the largest genome of all living things^36^. The observation that amoebae include the largest genome is based on qualitative data^36^. Amoebae genomes are characterized by various ploidy levels during their life cycle^37^; hence accurate estimation of genome size based on indirect techniques might not necessarily reflect the actual genome size. Determination of genome size should be accompanied by quantitative data using reliable techniques and sequence data.

Gene and GC contents vary in amoebae genomes. In general parasites belonging to the genus *Entamoeba* have smaller gene content^8^. The exception to this is *Entamoeba invadens*, which has more genes and GC content than its closest relatives (Table S6). *C. minus* is among species with larger gene models but is among species with lower GC content (Table S6). Gene and GC content do not seem to show evolutionary correlation in the currently available annotated genomes of amoebae (Table S6). Noticeably, *D. discoideum* (22.5%) and *A. castellanii* (58.4%) have the smallest and largest GC content percentages, respectively (Table S6). *C. minus* is closely related to *A. castellanii* (clade Centramoebida), however, there are stark differences in their GC and gene content (Table S6). The two major clades of Amoebozoa, Discosea and Evosea, are represented by more genomic data (both annotated and unannotated) than the third clade, Tubulinea (Table S6). A recent publication reported a draft genome of a tubulinid, *Vermamoeba vermiformis*^24^. Despite its small size, *V. vermiformis* is reported to have a relatively large genome (59.6 MB) and gene content (22,483 genes) compared to those available annotated genomes in the NCBI. Amoebozoa is a highly diverse group, more genome sequencing representing the three major clades is expected to unravel interesting patterns of genome evolution that would explain the great diversity observed in the supergroup.

### The nature of foreign gene acquisitions in Amoebozoa genomes

A considerable proportion of *C. minus* genome matches to foreign entities including bacteria, archaea and viruses (Fig. 1). Some of these foreign matching genes show evidence of trans-genomic trafficking through LGT (Table S3). The most dominant contribution of putative LGT in *C. minus* genome is bacteria. Similar large gene trafficking from bacteria are also reported in other amoebae^9,24^. Amoebae are major grazers of environmental bacteria. Permanent, obligatory and transient associations are known between bacteria and amoebae. Previous and our recent studies showed that large bacterial phyla including amoeba resisting bacteria and dangerous human pathogens are associated with amoebae^3,4,38^. These frequent and intimate encounters and associations might explain the observed large proportion of LGT events from bacteria in amoebae genomes. Bacteria-amoebae associations and LGT has significance on our understanding of bacterial pathogenesis and how pathogens can modulate host genome to escape host immunity. Several pathogenic bacteria are known to evade digestion and defense mechanisms of amoebae and other protists^39^. For this reason amoebae have been considered as a training ground for emerging pathogens^40^. Understanding the role of LGT in pathogenesis is critical to mitigate emerging human pathogens that can cause major public health concerns.

Unlike bacteria, the association of amoebae with archaea is poorly documented. Although archaea are hypothesized to be major contributors in the origination eukaryotic genome^41^, evidence of close associations and recent events of LGTs are few or rare in all studied amoebae genomes to date^9,24^.

Recent studies have reported an exciting discovery on association of protists with viruses^25,26,42^. Of particular interest is the association of giant viruses with amoebae^25,43^. Giant viruses have large filamentous capsids comparable to the size of bacteria. They also have large double-stranded DNA genomes encoding hundreds of genes^44^. The first giant virus, mimivirus, associated with amoeba was described from the genus *Acanthamoeba* (*A. polyphaga* mimivirus)^25^. Since their first discovery, a phylogenetically diverse groups of giant viruses have been described in amoebae and several other eukaryotes^42^. Interestingly, genes of giant virus origin have been discovered in the genomes of amoebae^9,24,42^ including *C. minus* (Table S3). These discoveries demonstrate that the association of giant viruses and amoebae must have a long-standing evolutionary history despite their recent discovery. Among the largest contributors of viral origin genes in *C. minus* is Pithovirus, the largest giant virus genus described to date^45^. Members of Mimivirus (Megaviricetes) originally described from *Acanthamoeba* are the largest putative LGT donors in the *C. minus* genome (Table S3). Similarly, Mimivirus is the largest contributor of putative LGT genes in the genomes of *Acanthamoeba*^9^ and *V. vermiformis*^24^. Although the majority of viral sequences in amoebae genomes come from giant viruses, a small fraction trace back their origin to double stranded DNA (dsDNA) bacteriophages and unclassified viruses (Table S3). Genes showing high similarity to Caudovirales, dsDNA bacteriophages, are also reported in *D. discoideum* and *E. histolytica*^9^ similar to *C. minus*. It is also interesting to note that all putative LGT donors in amoebae genome are dsDNA viruses. These findings clearly demonstrate that viruses play a role in the evolution of Amoebozoa. The nature and role of viruses in Amoebozoa evolution will become more evident as more genome data becomes available.

All putative *C. minus* gene models that we designated as LGT are found in scaffolds that are dominated with amoeba and eukaryotes genes. Expressed transcripts of some of these genes have been detected in transcriptome data collected from various life stages of *C. minus*. Statistical and phylogenetic analyses support the likelihood of some of these genes as putative LGTs. Some of these genes play a role in known biological processes (e.g., metabolism, cellular processes and signaling, information storage and processing), while some have unknown functions (Fig. S1). A closer examination of these putative LGT genes show that the majority have acquired introns (Table S3). The average number of introns across all coding genes in the *C. minus* genome is 4.7, slightly lower than *A. castellanii* (6.2) and higher than *V. vermiformis* (3.5)^24^. Putative LGTs in *C. minus* genome have more introns than the average genes transferred from bacteria having 6.2, those from viruses having 5.9, and those from archaea having 7.2 (Table S3). These results suggest that many of these transfer events were relatively ancient and that intronization of laterally transferred genes occurred at a rate higher than background perhaps due to either positive selection or at least a reduction in purifying selection.

### Genomic perspective of sex in Amoebozoa and *C. minus*

Members of the supergroup Amoebozoa display diverse life cycles involving sexual and asexual stages. Both molecular and cellular aspects of the diverse sexual life cycle observed in the Amoebozoa is poorly understood and documented only in a few lineages^6,28^. Recent genomic and transcriptomic work demonstrate that all amoebozoans examined possess genetic toolkit important for sexual development and genetic recombination^29^. However, the exact mechanism of sexual life cycles in the group remains elusive due to limited physical observations and genomic data.

Limited comparative genomic studies revealed that variation might exist in molecular mechanism of sexual cycle in amoebozoans^29^. As stated above *D. discoideum* and its closest relatives lack a recognizable *SPO11*, a gene important in initiation of meiotic recombination by introducing double strand breaks^27^. Variations in the numbers of *SPO11* paralogs also exist in the Amoebozoa. Similar to plants and some eukaryotes, *A. castellanii* possesses two copies of *SPO11*^30^. Three copies of *SPO11* are generally known in eukaryotes, two of which (*Spo11-1* and *Spo11-*2) are shown to be meiosis specific in *Arabidopsis*, while *Spo11-3* has a nonmeiotic role in plants^46,47^. *SPO11* has a complex evolutionary history involving lineages-specific duplications, losses and functional diversity in eukaryotes^30^.

Several eukaryotes including animals and fungi, possess only one copy of *SPO11* (a paralog of *Arabidopsis Spo11-1*), while some plants and a small number of protists that have been genomically surveyed, possess up to three copies of this gene ^30^. Examination of available Amoebozoa genomes reveals a spectrum of *SPO11* evolution in the supergroup. Similar to *A. castellanii, C. minus* possesses two copies of *SPO11* (*Spo11-1* and *Spo11-2*) (Fig. 4). In *Arabidopsis*, the two copies of *SPO11* are suggested to work as heterodimers^48^, while in eukaryotes with one copy of the gene, *SPO11* functions as a homodimer^30^. Transcriptome data analysis showed that several amoebae representing the three major clades of Amoebozoa possess *SPO11-2* (Fig. S2). Due to the incomplete nature of transcriptome data, the presence of the two copies of *SPO11* based on a phylogenetic analysis (see Fig. S2) can only be confirmed for *Gocevia fonbrunei* in amoebae with no genome data. Meiosis genes are expressed at low levels and can easily be missed in transcriptome data. Given the prevalence of *Spo11-2* in major groups of amoebozoans, it is likely that heterodimeric activity of *SPO11* in Amoebozoa is widespread (Fig. S2). An interesting *SPO11* evolution involving a lineage specific duplication is also observed in the parasitic genus *Entamoeba*^30^ (Figs. 4, S2,). Members of *Entamoeba* have duplicate copies of *Spo11-1* (*Spo11-1a* and *Spo11-1b*) in their genome adding complexity to the evolution of this gene within the supergroup. The variation of *SPO11* evolution in the Amoebozoa is indicative of the complex life cycle observed in the supergroup.

Some key genes involved in meiosis and karyogamy are not detectable in the *C. minus* genome. We were unable to detect the meiosis gene *HOP1* in *C. minus*. This was surprising as this gene is present in *A. castellanii* and other related amoebae^29,31^. Similarly, we were unable to detect *HAP2*, a gene involved in cellular fusion, and *GEX1/KAR5*, a gene involved in karyogamy, in the *C. minus* genome. This too was surprising given the extensive cellular and nuclear fusion behavior observed in *C. minus*

There are several possibilities to explain the apparent absence of these key meiosis and karyogamy genes. One is that the genes are not lost, but are instead not detectable by BLAST due to elevated evolutionary rates. If this were the case, the missing genes would have been labeled ORFans in our annotation. There are several genes considered ORFans that are upregulated during the fused stage of *C. minus* cells that could be considered candidates (Table S2). Among the upregulated ORFans, some genes contain functional domains involved in DNA damage sensing, replication and nuclear/chromosomal processes suggesting developmental roles (Table S2). Nevertheless, our finding clearly demonstrates that variation in sexual mechanism exist in the Amoebozoa. Future studies involving live experimentation, gene manipulation complemented with improved transcriptome and genome data will help elucidate the variations observed in the sexual mechanisms of amoebozoans at molecular level.

### Evidence of alternative splicing in a meiosis gene

Using genomic and transcriptomic data we find evidence of alternative splicing in one of the meiosis specific genes, *MER3*, in the *C. minus* genome. *MER3* is a conserved meiosis specific DNA helicase involved in ZMM-dependent crossover (class I cross-over) pathway^49^. Our previous work demonstrated that *MER3* plays a critical role in the sexual cycle of *C. minus*^20^. The analyzed *MER3* gene copy includes 6 domains (DEAD-Helicase-SEC63-DEAD-HELICASE-SEC63; Fig. 5). The repeated pairs of domains (e.g., the two DEAD domains) at this locus show significant primary sequence differences at both the nucleotide and amino acid levels. The transcript (mRNA), corresponding to *MER3*, found to be highly expressed in *C. minus* fused cells included only three of the six domains (i.e., the first two and the last, Fig. 5) indicating post-transcription processing. Alternative splicing of *MER3* has been also described in yeast^50^. It is likely that the *MER3* in *C. minus* undergoes similar splicing events as reported in fungi. This finding is a rare example of alternative splicing in Amoebozoa and provides insight into the complex regulation of the *MER3*. Our understanding of molecular processes in amoebozoans will improve as more high-quality genomes representing the diverse groups become available.

## Materials and Methods

### Genomic DNA collection of *Cochliopodium minus*

In this study, we sequenced the genome of *Cochliopodium pentatrifurcatum* ATCC© 30935TM. This species was found to be genetically identical to another strain of *Cochliopodium* (*C. minus* CCAP 1537/1A)^51^. These two isolates show distinct scale morphology but later it was found out that *Cochliopodium* species can express more than one type of scales during their life cycle. *C. pentatrifurcatum* have been synonymized to *C. minus* on the basis of this observation and genetic evidence^52^. In this study, *Cochliopodium minus* will be used to refer to the ATCC© 30935TM isolate. We used various approaches to collect genomic DNA from *C. minus*, which was grown in plastic petri dishes with bottled spring water (Deer Park®; Nestlé Corp. Glendale, CA) at room temperature supplemented with autoclaved grains of rice. First, genomic DNA from monoclonal whole culture was collected from actively growing cultures of *C. minus* maintained in our laboratory. Large number of cells from several petri dishes at maximum confluence were thoroughly washed with water to remove food bacteria. Cells were collected by gentle scraping in 15 ml tubes and centrifuged at 2000 rpm. Cell pellets were used to extract genomic DNA using MagAttract high-molecular-weight (HMW) DNA kit (Qiagen, MD), following the manufacturer’s instructions.

The second approach involved whole genome amplification (WGA) of single cells and nuclei pellets. For the single cells WGA approach, we picked and washed individual cells (∼100) using a mouth pipetting technique. For the nuclei extraction, monoclonal cells grown in 10 petri dishes were cleaned thoroughly and adherent cells were lysed by adding 6 ml lysis buffer (sodium phosphate buffer pH 7.4, 5 mM MgCl_2_, and 0.1% Triton-X 100) for 2 hours. The lysis step helps the release of nuclei into the cell culture (lysate). The lysate containing free nuclei were collected and centrifuged for 10 minutes at 500 rpm at room temperature. The nuclei pellet was then re-suspended in 0.5 ml of lysis buffer and transferred on top of 12 ml sucrose cushion. The sucrose cushion aids in separation of nuclei by trapping small particles (e.g., bacteria) or light weighted lysates (cell parts) through centrifugation. The lysate and sucrose cushion mixture were centrifuged at 3200 rpm for 20 minutes at room temperature. The pellet from this step was resuspended in 1 ml of lysis buffer and centrifuged again at 10,000 rpm for 1 minute at room temperature. The purified nuclei pellets were collected after carefully removing the supernatant. Nuclei pellets and single cells were used to perform WGA using REPLI-g Advanced DNA single cell kit (QIAGEN; Cat No./ID: 150363) according to the manufacturer’s protocol. Amplified DNA was quantified using Qubit assay with the dsDNA broad range kit (Life technologies, Carlsbad, CA, USA).

### Genomic DNA library preparation and sequencing

We applied three different sequencing strategies: Illumina short reads, 10x genomics linked reads, and Oxford Nanopore long read sequencing. For Illumina short read sequencing, we sent a nucleus pellet amplified gDNA sample to GENEWIZ (South Plainfield, NJ) for library preparation and sequencing. Sequencing was performed using an Illumina HiSeq instrument, which generated pair-end, high-output mode with 150 bp reads. We also applied a linked-read sequencing strategy using the 10X genomics platform. We sent whole culture gDNA to the Yale Center for Genomic Analysis for library preparation and 10X sequencing. For long read sequencing, we used the Oxford Nanopore technology (ONT) (Oxford Nanopore Technologies Ltd., United Kingdom) with the MinION device using the SQK-RAD004 kit. We constructed the library from 400 µg of amplified nuclear or single cells gDNA and this library mix was added to the flow cell using the SpotON port.

### *De novo* Genome assembly

A set of step-by-step commands outlining the genome assembly process are included in Supplementary file-1. Software versions, citations, and links to software repositories are provided in this supplementary document. Direct assemblies of long read and linked read data produced assemblies with a mix of independently assembled allelic contigs (haplotigs) interspersed with collapsed haplotypes and were therefore not satisfactory despite high contiguity in these assemblies. We implemented the following hybrid approach to circumvent this problem.

We generated three sets of genomic data using different technologies for each. This included: (*1*) 333 million Illumina paired end reads after trimming, (*2*) 256,923 MinION long reads, and (*3*) 267 million 10X Genomics chromium linked reads. We trimmed adapters from Illumina paired end reads using BBDUK. We assembled Nanopore reads using Canu (Supplementary file-1). We assembled the adapter-trimmed Illumina reads using the SPAdes assembler and provided a FASTA file with Canu contigs that were 10 Kb and longer as a set of trusted contigs to the Spades assembler (Supplementary file-1). We next ran Redundans on the resulting SPAdes assembly to reduce heterozygous regions of the genome that were represented more than once to a single representative and to remove very short contigs.

We assembled the 10X Genomics linked reads using SuperNova (Supplementary file-1). We then generated artificial mate pairs using MateMaker with insert sizes ranging from 200–50,000 bp. We then used SSPACE to scaffold our SPAdes assembly with these mate pair libraries. We divided the resulting assembly at scaffolding points that produced gaps greater than 10Kb.

### Contaminant removal

We performed the following operations to filter out contamination from food bacteria, symbionts (viruses and archaea) and other environmental contaminants. First, we used Basic Local Alignment Search Tools (BLAST 2.10.0+) to identify single best BLASTN match for each assembled scaffold using the locally installed NCBI non-redundant nucleotide (nt) database. Hits with >90% identity and >90% query coverage to bacterial, archaeal or virus sequences were removed. We used the remaining scaffolds for gene prediction (see below). We then searched the NCBI non-redundant protein database (nr) using BLASTP. We inspected each scaffold with a significant BLASTP hit to a potential contaminant and removed any scaffold where most of the genes on that scaffold had best BLAST hits to the same bacteria, archaea, virus or non-amoeboid eukaryote were excluded. We assessed the decontaminated scaffolds with BUSCO^53^ and transcriptome data to ensure that the approach did not remove amoeba genes.

### Gene prediction and functional annotation

A set of step-by-step commands outlining the genome annotation process are included in Supplementary file-1. We used BRAKER2 in combination with RNA sequencing data of *C. minus*^11,20^ and the protein sequences of a published genome of a closely related amoeba, *Acanthamoeba castellanii* to annotate the *C. minus* genome assembly described above. We aligned *C. minus* RNA-seq data to the genome assembly using STAR (Supplementary file-1).

We classified likely homologs in and associated Clusters of Orthologous Groups (COGs) categories with our predicted sequences using the EggNOG-mapper implemented in OmicsBox v.2.0.29. We used BLASTP to search gene models against the nr database. Genes that had no hits to nr were classified as ORFans. These genes were further investigated to retrieve their putative functions based on domain search in Hmmer web server v. 2.41.1 against the reference proteome database with default parameters (https://www.ebi.ac.uk/Tools/hmmer/). Genes that had significant domain hits were further analyzed to determine their roles in different life stages of the amoeba through a differential gene expression (DGE) analysis. The DGE analysis was modified based on methodology described in Tekle et al.^20^, where the genome data is used to map RNA-seq reads. We assessed the completeness of the draft genome, with the gene models, using Benchmarking Universal Single-Copy Orthologs v.5^53^ against the eukaryotic database of 250 genes via the web assessment tool gVolante (https://gvolante.riken.jp/).

### Lateral gene transfer (LGT) and Sex genes analyses

We compared the protein models against the nr database using BLASTP with a threshold e-value of 1×10^−10^. We retrieved the taxonomic affiliation of hits from NCBI taxonomy using a Perl script in the KronaTools v2.7.1 package (https://github.com/marbl/Krona/releases/tag/v2.7.1). We then carried out an Alien Index (AI) analysis to identify likely sources of lateral gene transfer. We used Alien Index v2.1 (https://github.com/josephryan/alien_index) with default parameters. We considered protein models with alien index scores ≥ 45 as foreign, 0□≤□AI□≤□45 as indeterminate, and less than 0 as amoeba genes. We also searched previously identified LGTs from related amoebae (*E. histolytica, E. dispar, A. castellanii* and *D. discoideum*)^9^ in the genome of *C. minus*. From this analysis, putative LGTs that had significant hits to *C. minus* genes and were not recovered with our Alien Index analysis were added to the list of *C. minus* putative LGTs. Selected putative LGTs representing bacteria, archaea and viruses were further examined by building phylogenetic trees in IQ-Tree using the automatic model selection option and 1000 ultrafast bootstrap replicates^54^. Protein sequences for this analysis were aligned in AliView^55^ complied using BLASTp against respective domain and various representative eukaryotic groups with best blast matches for each putative LGT.

We performed gene inventory analyses of more than 90 genes including meiosis specific and sex related genes (fusion and karyogamy) using the draft genome of *C. minus* as in Wood et al.^21^. The domain search of selected genes was performed using phmmer search as implemented on the web server, https://www.ebi.ac.uk/Tools/hmmer/search/phmmer (Supplementary file-1). To ensure that *C. minus* homolog position was compatible with other isoforms from other organisms, we perform phylogenic analyses in IQ-Tree^54^ as described above to select the correct orthologs.

## Acknowledgments

This work is supported by the National Science Foundation EiR (1831958) and National Institutes of Health (1R15GM116103-02) to YIT. This work was also supported by the National Science Foundation under grant number 1542597 to JFR. We would like to thank James T. Melton III, Fiona Wood, Stephen Kioko and Maya Blasingame for technical assistance during data collection and analysis.

## Author contributions

YIT conceived the project, led writing manuscript and helped design experiments and analyses. FW and HT collected data, conducted analyses, and contributed to writing and editing of the manuscript. DH helped with genome assembly analyses, general discussion, and writing. JFR helped design genome assembly pipeline and edited the manuscript. All authors have read and approved the manuscript.

## Supplementary Materials Captions

**Figure S1**. Functional categories based on Cluster Orthologous Groups (COGs) database of putative LGTs in bacteria, viruses and archaea.

**Figure S2**. Phylogenetic reconstructions of SPO11 paralogs including more amoebozoan taxa using transcriptome data. Transcript from some amoebae group with *SPO11-2* supporting the prevalence of *SPO11-2* in other amoebozoan lineages. Among amoebae with transcriptome data, *Gocevia fonbrunei* has both copies. Note the presence of in-paralog (lineage specific duplication) of *SPO11-1* in the parasitic genus *Entamoeba*. Clade supports at nodes are ML IQ-TREE 1000 ultrafast bootstrap values. All branches are drawn to scale.

**Figure S3**. Phylogenetic reconstructions of MER3 paralogs. Two copies of MER3 genes from the *Cochliopodium minus* draft genome are placed into two separate poorly supported clades. The copy, jg36575.t1, displaying alternative splicing and that is highly expressed in fused cells of *C. minus* fall with canonical copy of MER3 found in other eukaryotes. Clade supports at nodes are ML IQ-TREE 1000 ultrafast bootstrap values. All branches are drawn to scale.

**Table S1**. Distribution of the gene models of *Cochliopodium minus* draft genome in categories of clusters of orthologous groups of proteins (COGs).

**Table S2**. Differential Gene Expression of ORFan genes using PFAM scan between fused *versus* unfused cells of *Cochliopodium minus*.

**Table S3**. Putative LGT-derived genes in *Cochliopodium minus* genome with Alien Index above threshold (>45) scores.

**Table S4**. Virus matching gene models of *Cochliopodium minus* genome.

**Table S5**. Tyrosine Kinase genes-based om InterProScan analysis of the *Cochliopodium minus* genome.

**Table S6**. Genome size, gene content and GC content of amoebae that are annotated and unannotated genome available in the NCBI.

